# Ethylene signaling is essential for mycorrhiza-induced resistance against chewing herbivores in tomato

**DOI:** 10.1101/2024.06.13.598897

**Authors:** Javier Lidoy, Javier Rivero, Živa Ramšak, Marko Petek, Maja Križnik, Victor Flors, Juan A. Lopez-Raez, Ainhoa Martinez-Medina, Kristina Gruden, Maria J. Pozo

## Abstract

Root colonization by certain beneficial microbes can prime plant defenses aboveground, modifying plant responses to potential attackers. Arbuscular mycorrhizal (AM) fungi establish mutualistic symbiosis with most plant species, usually enhancing plant resistance to biotic stresses, leading to Mycorrhiza-Induced Resistance (MIR). Still, our knowledge of the complex molecular regulation leading to MIR is very limited. Here we show that the AM fungus *Funneliformis mosseae* protects tomato plants against two different chewing herbivores, *Spodoptera exigua* and *Manduca sexta*, and we explore the underlying molecular mechanism.

We explore the impact of AM symbiosis on the plant response to the herbivores through genome-wide transcriptional profiling, followed by bioinformatics network analyses and functional bioassays. Herbivore-triggered JA-regulated defenses were primed in leaves of mycorrhizal plants, while ET biosynthesis and signaling was also higher both before and after herbivory. We hypothesized that fine-tuned ET signaling is required for the primed defensive response leading to MIR in mycorrhizal plants. We followed analytical, functional, and genetic approaches to test this hypothesis and get mechanistic insights into the ET signaling in MIR.

ET is a complex regulator of plant responses to stress, and although ET is generally considered a negative regulator of plant defenses against herbivory, tomato lines deficient in ET synthesis or perception could not develop MIR against either herbivore. Thus, we demonstrate that hormone crosstalk is central to the priming of plant immunity by beneficial microbes, with ET fine-tuning being essential for the primed JA biosynthesis and boosted defenses leading to MIR in tomato.

## 1. Introduction

Upon herbivory, plants activate an extensive arsenal of defensive mechanisms including toxic, anti-digestive, and repellent compounds (Broadway & Duffey, 1988; Duffey & Stout, 1996; Erb & Reymond, 2019). The jasmonate (JA) signaling pathway plays a central role in the transcriptional reorganization and the induction of defenses in the process (Wasternack & Hause, 2013; Erb & Reymond, 2019; Wang *et al*., 2019). Jasmonates are oxylipins synthesized from linolenic acid in the chloroplast membranes into 12-oxo-phytodienoic acid (OPDA). Subsequently OPDA is oxidized in the peroxisome to jasmonic acid, which can be further metabolized to various derivatives, including conjugates as jasmonoyl-L-isoleucine (JA-Ile) which binds to the CORONATINE INSENSITIVE1 (COI1) receptor that triggers the transcriptional regulation of the defense response (Yan *et al*., 2009). Other phytohormones contribute to shaping the defense response through crosstalk with other hormone signaling pathways, adjusting the plant response to the context perceived (Erb *et al*., 2012; Berens *et al*., 2017; Aerts *et al*., 2021). Among them, ethylene (ET) is a gaseous phytohormone produced from the oxidation of its precursor, 1-Aminocyclopropane-1-Carboxylate (ACC) by ACC oxidases (ACO) (Yang & Hoffman, 1984). This phytohormone is involved in multiple processes from plant growth, development and defense, fruit ripening and abscission, senescence and tolerance to diverse abiotic stresses (Dubois *et al*., 2018). It also contributes to the regulation of plant defense and interactions with other organisms through the modulation of JA dependent responses (Broekgaarden *et al*., 2015). In arabidopsis, two main branches in JA regulated defenses are described, the JA/ET branch, coordinating defenses against necrotrophic pathogens, and the JA/ABA coordinated branch, effective against chewing herbivores, and they are believed to be mutually antagonistic (Verhage *et al*., 2011; Kazan & Manners, 2013; Zhang *et al.,* 2018). Thus, ET is considered antagonist to JA-dependent defenses against herbivory. However, recent findings point to a more complex role for ET, as novel connections between these two pathways have been identified (Ma *et al.,* 2020; Hu *et al.,* 2021), but the relevance of JA-ET crosstalk in shaping antiherbivory responses is yet to be uncovered.

Under natural conditions, plants interact simultaneously with multiple organisms, and this leads to a complex finetuning of the plant responses, commonly mediated by hormone crosstalk, that shape the final outcome of the multiway interactions (Gruden *et al*., 2020). Most of the molecular mechanisms that regulate two-way interactions between plants and arthropods are also activated in plant-microbe interactions, but in multiway interactions between plants, microbes and insects the plant response becomes more complex: additional pathways can be activated, and changes in the intensity and timing of the responses are common (Gruden *et al*., 2020). In this regard, plant associated microorganisms, including arbuscular mycorrhizal (AM) fungi, can alter plant responses to attackers aboveground, frequently leading to induced plant systemic resistance to diverse pathogens and pests (Pieterse *et al*., 2014; Pozo *et al*., 2021).

Over 70% of all vascular plants can establish a mutualistic symbiosis with AM fungi (Brundrett & Tedersoo, 2018). The AM symbiosis confers different benefits to the plant, from better mineral nutrition to better tolerance to biotic and abiotic stresses (Smith & Read, 2008), increasing plant resilience to cope with environmental challenges (Rivero *et al*., 2018). It can induce resistance to a broad range of pathogens and pests (Jung *et al.,* 2012; Song *et al.,* 2013, 2015; Sanmartín *et al.,* 2020b; Rivero *et al.,* 2021, Fiorilli *et al.,* 2024). The impact on pest feeding and performance may result from changes in the plant nutritional value and chemistry, hormone signaling and defense responses (Rivero *et al.,* 2021; Dejana *et al.,* 2022; Ramírez-Serrano *et al.,* 2022, Manresa-Grao *et al.,* 2022). Mycorrhiza-induced resistance (MIR) to pathogens and pests commonly rely on the priming of plant defenses (Pozo & Azcón-Aguilar, 2007; Jung *et al*., 2012). Priming of plant defenses (or immune priming) is a cost-efficient, adaptive defense strategy, in which preconditioned tissues are able to activate more efficiently plant immune responses upon challenge - usually leading to faster or stronger defense responses- (Conrath *et al*., 2006, 2015; Martinez-Medina *et al*., 2016; Mauch-Mani *et al*., 2017). Different experimental evidence points to a prominent role of JA signaling in the defense priming displayed in mycorrhizal plants (Jung *et al*., 2012; Song *et al*., 2013; Mora-Romero *et al*., 2014; Sanmartín *et al*., 2020b). However, little is known about the contribution of other hormone signaling pathways to the regulation of mycorrhiza triggered JA-defense priming of MIR (Fiorilli *et al.,* 2024).

Here, we aimed to explore the molecular mechanisms regulating MIR in response to chewing herbivores. We first performed a genome-wide transcriptional profiling of the interactions between tomato, AM fungi and two different chewing herbivores. Mycorrhizal colonization led to primed JA-regulated responses to the herbivory challenge, but also to a differential regulation of ET signaling before and after herbivory. We found enhanced ET production in mycorrhizal plants, and network analysis pinpointed relevant connections between ET signaling and JA biosynthesis. We hypothesized that ET signaling plays a relevant role in the finetuned JA regulated responses of mycorrhizal plants to herbivory. To test the hypothesis, we used tomato lines deficient in ET synthesis and signaling, and found that the mycorrhiza associated primed responses were lost in these lines, and they were unable to display MIR against the herbivores. Our results show that differential regulation of ET signaling in mycorrhizal plants is essential for boosting JA production and JA dependent defenses against both herbivores in tomato, and it is, therefore, a key component of MIR.

## 2. Materials and Methods

### 2.1. Biological material and growing conditions

*Funneliformis mosseae* (T.H. Nicolson & Gerd.) C. Walker & A. Schüßler (BEG12, International Bank of Glomeromycota, https://www.i-beg.eu/cultures/BEG12.htm) is maintained in a pot culture of *Trifolium repens* L. and *Sorghum vulgare* Pers. in a substrate consisting of vermiculite:sepiolite (1:1, v/v) under greenhouse conditions. Tomato seeds were surface sterilized in 4% sodium hypochlorite for 10 min, washed with water and incubated in plastic trays containing sterile vermiculite at 25–27 °C, 16/8 h (day/night) and 65–70 % RH. After 10 days, plantlets were transplanted to 350 mL pots with sand:vermiculite (1:1, v/v). Mycorrhizal treatments consisted of plants inoculated with 10 % (v/v) of *F. mosseae* inoculum containing colonized root fragments, spores and mycelia. Plants were randomly distributed and kept under greenhouse conditions (25–27 °C, 16 h/8 h (day/night), 65–70 % RH). The fertigation schedule included watering with half-strength Hoagland solution (Hoagland & Arnon, 1938) once a week containing 25% of standard phosphorus. *S. exigua* Hübner (Lepidoptera: Noctuidae) eggs were obtained from the iDiv (Germany) for experiment 1, and from Universitat de València (Spain) for experiment 2. *S. exigua* larvae were reared on artificial diet (Hoffman & Lawson, 1964) and maintained at 24 °C. *M. sexta* L. (Lepidoptera: Sphingidae) eggs were obtained from iDiv (Germany) for experiment 1 and from Universität Osnabrück (Germany) for experiment 2. Eggs were incubated at 26 °C and larvae were reared on detached tomato leaflets.

### 2.2. Transcriptional profiling experiment

Surface sterilized tomato seeds (*Solanum lycopersicum* L. cv. Moneymaker) were used for the transcriptional profiling experiment. The experiment consisted of 6 treatments: Non-mycorrhizal control plants without herbivory (Nm), Non mycorrhizal plants challenged with *S. exigua* (NmSe), Non mycorrhizal plants challenged with *M. sexta* (NmMs), *F. mosseae* inoculated control plants without herbivory (Fm), *F. mosseae* inoculated plants challenged with *S. exigua* (FmSe) and *F. mosseae* inoculated plants challenged with *M. sexta* (FmMs). Each treatment consisted of 6 independent plants as biological replicates. After 5 weeks, a well-established mycorrhizal colonization was confirmed, and the herbivory assays were performed. Three 3^rd^ instar *S. exigua* larvae or two neonate *M. sexta* larvae were placed on the three apical leaflets of the third true leaf inside a clip cage (30 mm Ø). After 24 h, the larvae were removed, and the leaflets were frozen immediately in liquid nitrogen and stored at −80 °C. The extent of damage was assessed to confirm herbivory and the damage between herbivores was similar.

### 2.3. Functional analysis of MIR in ethylene deficient lines

Seeds of *S. lycopersicum* wild-type UC82B, ET-deficient line ACD (Klee *et al*., 1991) and ET-insensitive mutant never ripe (Nr) (Wilkinson *et al*., 1995) were surface sterilized and germinated as described above. After 8 weeks the herbivory assays were performed on mycorrhizal and non-mycorrhizal plants. Four 3^rd^ instar *S. exigua* larvae or three neonate *M. sexta* larvae were placed on the third fully expanded leaf inside an entomological bag. For the *S. exigua* bioassay, each treatment consisted of 7 independent plants, and 4 larvae were used per plant (a total of 28 larvae per treatment). For the *M. sexta* bioassay, 10 independent plants were used per treatment, and 3 larvae were used per plant (30 larvae per treatment). Larval mortality and pupation were monitored every 48 h and *M. sexta* weight was determined at 9 days post infestation.

### 2.4. Mycorrhizal quantification

As described in García et al. (2020), root samples were cleared and digested in 10% KOH (w/v) for 2 days at RT (18 – 23°C). Then, root samples were rinsed thoroughly with tap water and acidified with 2% (v/v) acetic acid solution. Fungal root structures were stained with a 5% (v/v) black ink (Lamy, Germany) and 2% acetic acid solution for 24 h at RT (Vierheilig *et al*., 2005). Ink solution was washed with tap water. Mycorrhizal colonization was determined by the gridline intersection method (Giovannetti & Mosse, 1980) using a Nikon SMZ1000 stereomicroscope.

### 2.5. Ethylene emission quantification by gas chromatography

One detached leaflet of each tomato plant was placed into a 20 mL glass vial containing a sterile filter paper soaked in 200 µL of sterile distilled water to avoid dehydration. The vials were left uncovered for 30 min to avoid the detection of the ethylene released as a result of the scalpel-induced wounds. After this time the non-herbivory treatments vials were sealed. For the herbivory treatments, we placed inside the vial one larvae of the corresponding herbivore and immediately after the vials were sealed. Vials were maintained at 23 °C under a 18 h photoperiod. 1 mL from each vial was withdrawn with a syringe and the area of the ethylene peak was analyzed in a gas chromatograph with a flame-ionization detector (GC-FID; Hewlett Packard 5890). ET emission by the herbivores were determined to be negligible by analyzing vials containing only larvae.

### 2.6. RNA-Seq transcriptional analysis

For RNA-Seq analysis, three apical leaflets of the third true leaf contained within the clip cage were harvested and immediately flash frozen in liquid nitrogen. Three biological replicates per treatment were used, each consisting of pooled material from two plants. Samples were grinded in liquid nitrogen. Total RNA was extracted with the RNeasy Plant Mini Kit (Qiagen, Germany) following the manufacturer’s instructions. The quality, quantity and size of extracted RNA was determined with a Bioanalyzer (Agilent, USA) and Nanodrop (ThermoFisher Scientific, USA). All samples were of good RNA quality (RIN > 8, A260/A280 > 1.8 and A260/A230 > 2). TruSeq stranded RNA-Seq library preparation and paired-end sequencing on the Illumina NovaSeq 6000 platform were performed by Macrogen (S. Korea). Quality control of sequencing reads was performed using FastQC (Andrews *et al*., 2010). Sequences were mapped to the tomato genome version SL4.0 using STAR v2.7.2b (Dobin *et al*., 2013) and the ITAG 4.0 annotation (Hosmani *et al*., 2019). Differential expression analysis was performed in R using the DESeq2 package v1.26.0 (Love *et al.,* 2014). Prior to statistical testing, genes not having at least 50 counts in at least three samples were excluded. Genes with FDR-adjusted p-value < 0.05 were considered significantly differentially expressed (DEG). Gene set enrichment analysis (GSEA) was performed with TMM normalized count values using the GSEA tool v4.0.3 (Subramanian *et al*., 2005) and gene sets based on GoMapMan (Ramšak *et al*., 2014) BINs. Gene sets with FDR-adjusted q-values < 0.05 were considered significantly enriched in up- or down-regulated genes. For easier visualization of the enriched gene sets, they were selected and organized in functional supergroups (Supplemental Table S1).

### 2.7. Network analyses

Network analyses were performed on a specifically generated knowledge network of *S. lycopersicum*. First, the *Arabidopsis thaliana* large comprehensive knowledge network (Ramšak *et al*., 2018), containing high-quality relations (protein-protein binding, protein-DNA binding, miRNA-transcript targets) between Arabidopsis genes was translated to tomato using PLAZA orthologues (Proost *et al*., 2015). Additionally, a tomato specific network of miRNA - transcript targets was generated using psRNATarget (Dai *et al*., 2018) for tomato miRNAs present in miRBase v22 (Kozomara *et al.,* 2019) and merged with the translated comprehensive knowledge network. A subnetwork with genes that were differentially expressed in at least one of the RNA-Seq pipeline contrasts (FDR adjusted p-value <= 0.05) was extracted in the next step. Shortest path searches were performed using EIN3/EIL1 transcription factors as starting nodes (Solyc01g009170, Solyc01g096810, Solyc06g073720, Solyc06g073730) and genes related to JA biosynthesis for the end nodes (LOX: Solyc01g099190, Solyc03g006540, Solyc03g122340, Solyc05g014790, Solyc08g014000; AOC: Solyc02g085730; AOS: Solyc04g079730, Solyc10g007960, Solyc11g069800; OPR: Solyc07g007870, Solyc10g086220, Solyc11g013170). Network analyses were performed in R using the igraph package v1.2.8 (Csárdi & Nepusz, 2006) and results visualized in Cytoscape (Shannon *et al*., 2003).

### 2.8. Analysis of gene expression by qPCR

Total RNA was extracted from leaves using TRIsure™ (Bioline, USA) and treated with DNase I (NZYtech, Portugal). Later, RNA was purified and concentrated using RNA Clean & Concentrator-5 column kit (Zymo Research, USA). First-strand cDNA was synthesized from 1 µg of purified total RNA using PrimeScript RT Master Mix (TaKara, Japan) according to the manufacturer’s instructions. qPCR reactions and relative quantification of specific mRNA levels were performed with a StepOnePlus™ Real-Time PCR System (Applied Biosystems, USA) and the gene-specific primers described in Supplemental Table S2. Expression values were normalized using the reference gene *SlEF-1α* (López-Ráez *et al*., 2010), encoding the tomato translation elongation factor-1α using the comparative 2^-ΔΔCt^ method (Livak & Schmittgen, 2001). For more robust statistical analysis of the differences in expression levels, 6 independent biological replicates per treatment were analyzed.

### 2.9. LAP enzymatic activity

Powdered leaf tissue was mixed 1:18 (w:v) with protein extraction buffer (50 mM Tris-HCl [pH 8] and 0.5 mM MnCl_2_). Samples were centrifuged at 14000 rpm for 10 min at 4 °C and the supernatant was collected. This process was repeated twice. For the LAP enzymatic activity, a stock solution of L-leucine-*p*-nitroanilide (LpNA; Sigma-Aldrich, Germany) was prepared in absolute ethanol. The reaction mixture contained 200 µL of 3 mM LpNA (in 50 mM Tris-HCl [pH 8] and 0.5 mM MnCl_2_) and 40 µL of the sample protein supernatant.

The reaction was incubated in a 96-well plate at 37 °C for 20 min. The absorbance was measured at 410 nm (Chao *et al*., 2000).

### 2.10. Phytohormone analysis

Freeze dried powdered leaf material was used for hormonal analysis as described by Sánchez-Bel et al. (2018) with small changes. 30 mg of dry material was homogenized with 1 ml MeOH:H2O with 0.01% of HCOOH containing a pool of a mixture of internal standards of jasmonic-2,4,4d_3_-(acetyl-2,2-d_2_) acid (Sigma-Aldrich) for both JA and OPDA and own synthesized JA-Ile-d_6_ quantification up to a final concentration in sample of 3 µg.l^-1^. Samples were ground in cold and centrifuged at 15000 rpm for 15′. The pH of the supernatant was reduced to 2.5–2.7 with acetic acid and the extraction was partitioned twice against diethyl ether. The organic phase was recovered and evaporated in a speedvac centrifuge. Samples were resuspended in 1 ml of H2O/MeOH (90:10) with 0.01% of HCOOH up to a final concentration of internal standards of 10 ng ml^-1^. The chromatography was performed using a UPLC Kinetex C18 analytical column with a 5 μm particle size, 2.1 100 mm (Phenomenex). Samples were injected onto an Acquity ultraperformance liquid chromatography system (UPLC; Waters, Mildford, MA, USA), which was interfaced with a triple quadrupole mass spectrometer (TSD, Waters, Manchester, UK). Quantification was performed by using Masslynx 4.2 software.

### 2.11. Statistical analyses

Besides the methods and software for RNA-Seq transcriptional analysis described above, statistical analyses were performed with unpaired t-test analysis using Statgraphics Plus 3.1. Comparison between treatments of larval mortality and pupation was performed using the Log-Rank test (Mantel-Cox) with the “survival” and “survminer” packages in R. PCAs were performed using Metaboanalyst software.

## 3. Results

### 3.1. Mycorrhizal symbiosis impacts transcriptional regulation

We previously showed that *F. mosseae* induced resistance in tomato against *S. exigua* by priming the accumulation of antiherbivore compounds (Rivero *et al*., 2021). To explore the molecular processes underlying the impact of *F. mosseae* on insect performance, we conducted an untargeted transcriptomic study in tomato leaves. We compared the full transcriptional profile of leaves from non-mycorrhizal (Nm) and mycorrhizal plants colonized by *F. mosseae* (Fm) without challenge, or subjected to herbivory by the generalist *S. exigua* (NmSe, FmSe) and the specialist chewing herbivore *M. sexta* (NmMs, FmMs). Principal Component Analysis (PCA) was performed on the RNA-Seq data (Fig. 1A). The first two principal components explained 75.5% of the total variance. Herbivory had a strong impact on the transcriptome, with a clear separation from non-herbivory treatments in the PCA plot. This separation was mostly explained by PC 1, which accounts for 63.8% of the total variance (Fig. 1A). In contrast to the strong effect of herbivory on the leaf transcriptome profile, mycorrhizal colonization itself did not show a significant effect (Nm and Fm; Fig. 1A). However, when focusing only on herbivore challenged plants, a new multivariate analysis revealed a significant impact of mycorrhization in the transcriptomic profile under herbivory, as illustrated by the separation, mostly explained by PC1, between the non-mycorrhizal (Nm) and mycorrhizal (Fm) plants upon challenge with any of the herbivores (FmSe and FmMs vs NmSe and NmMs; Fig. 1B), pointing to a differential plant response to the herbivore upon mycorrhization.

**Figure 1.**
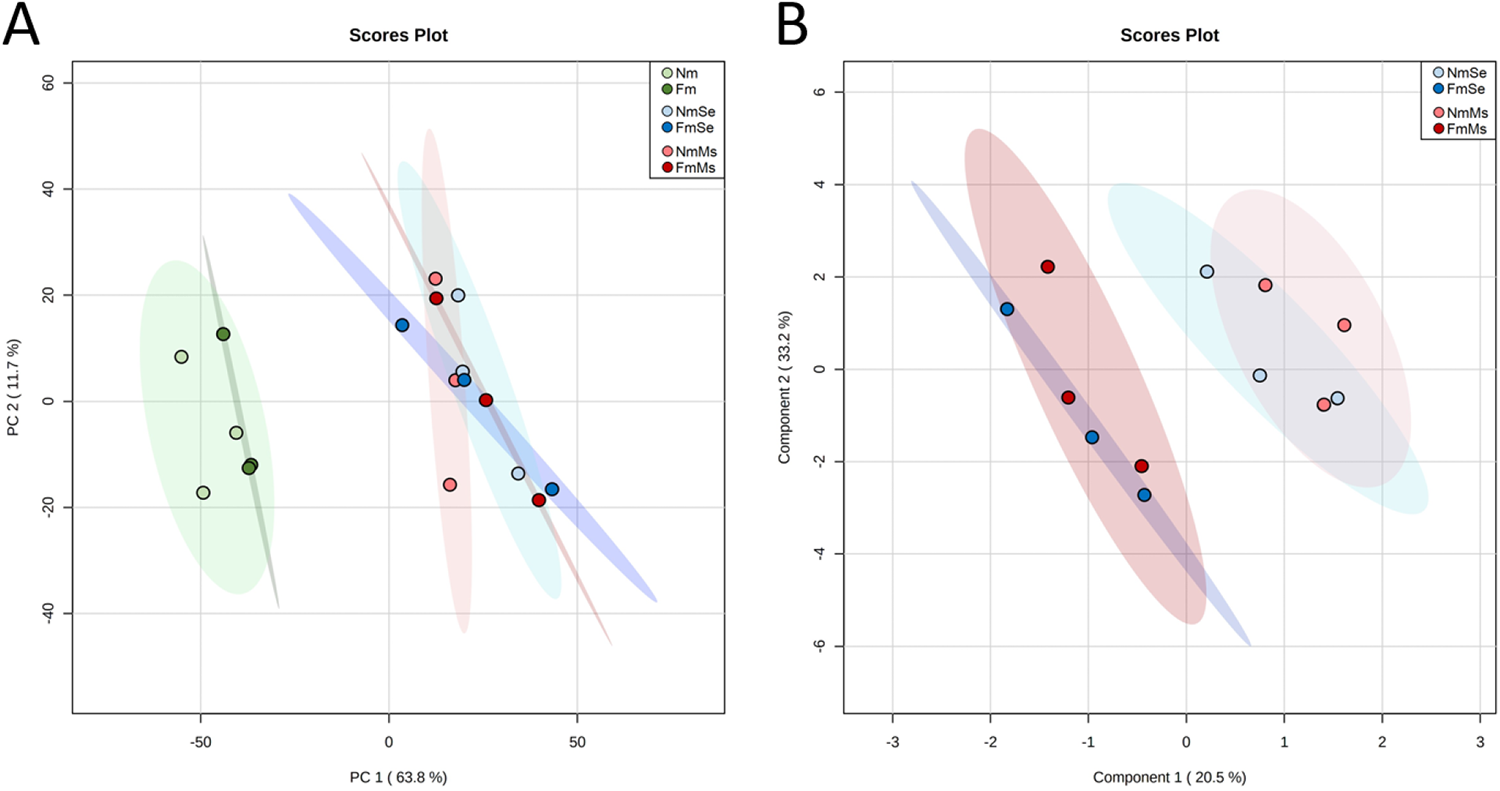
Overview of the impact of the different treatments on the tomato leaf transcriptome. Tomato leaves from uninfested non-mycorrhizal (Nm) and *F. mosseae* mycorrhizal (Fm) plants, or subjected to 24h of *S. exigua* (NmSe, FmSe) or to *M. sexta* (NmMs, FmMs) herbivory. (A) Overall transcriptomic PCA plot for all treatments. (B) Transcriptomic PLS-DA 2-D Scores plot between herbivory treatments only. The percentage of variance explained by the principal component is shown in brackets.

### 3.2. Mycorrhizal colonization modulate key regulatory pathways

In the absence of herbivory, only 57 differentially expressed genes (DEGs) (FDR <0.05, Supplemental Table S3 and S4) were identified in Fm plants as compared to the Nm controls. We explored the transcriptional changes through gene set enrichment analysis (GSEA), an analytical method focusing on the regulation of gene groups sharing common biological functions (Subramanian *et al.,* 2005). The GSEA revealed the modulation of several important cell processes in Fm plants (Fm; Fig. 2). Besides changes in cell division and structure (especially in “cell cycle” and “cell wall structure”) most changes were found in signaling related pathways (“sugar and nutrient signaling”, “receptor kinases signaling”), gene sets related to secondary metabolism (increase in “stress biotic receptors’’ and “glycosyl and glucoroyl transferases”), and pathways related to the plant stress responses (“DNA chromatin structure”, “protein metabolism”, “RNA regulation”). Hormone metabolism was also differentially regulated by *F. mosseae* colonization, as revealed by the enrichment of genes related to the JA and ET pathways (Supplemental Tables S6 and S7). Noteworthy, a high number of transcription factors, particularly related to ethylene (ET) signaling, were observed among the Fm regulated genes (Supplemental Table S5).

Herbivory had a strong impact on the leaf transcriptome, with 6172 and 4534 DEGs in leaves challenged by the generalist *S. exigua* and the specialist *M. sexta,* respectively (Supplemental Tables S3 and S4). When comparing directly plants challenged by the different herbivores (NmSe vs NmMs), no DEGs were found (Supplemental Table S5), suggesting common plant responses to herbivory, probably varying in intensity – the specialist leading to lower changes in the host-. In fact, the GSEA revealed that the general response to both herbivores is conserved, with most functional classes-related to both primary and secondary metabolism-regulated similarly (Fig. 2). Herbivory challenge repressed gene expression related to photosynthesis and impacted synthesis and degradation of amino acids and protein metabolism. Both herbivores activated secondary metabolism, inducing the synthesis of phenylpropanoids and lignins as well as PR proteins with antiherbivory functions (mainly proteinase inhibitors, proteases, polyphenol oxidases) (Fig. 2; Supplemental Table S4). Herbivory also impacted hormone metabolism, mostly by enriching the JA metabolism related gene set (Fig. 2; Supplemental Table S6). Globally, the impact of the specialist *M. sexta* on the primary metabolism was lower that of the generalist *S. exigua,* with a lower repression of photosynthesis and protein synthesis related genes (Supplemental Fig. S1).

When focusing on the mycorrhizal effect, we found that the core responses to herbivory were overall similar in mycorrhizal and non-mycorrhizal plants. As in non-mycorrhizal treatments, mycorrhizal plants responded to herbivory with an induced expression of genes related to secondary metabolism and JA-dependent PR proteins involved in defense against chewing herbivores (Fig. 2) and repression of the primary metabolism, although this repression was stronger in mycorrhizal plants than in non-mycorrhizal ones (FmSe, FmMs; Supplemental Fig. S2). Despite the similar responses to herbivory, we found specific differences between herbivory-challenged mycorrhizal and non-mycorrhizal plants related to cell wall synthesis and modification and related to hormone metabolism. While the activation of the JA pathway upon herbivory was common in both Nm and Fm plants, the ET and the ABA metabolism were up-regulated by herbivory only in mycorrhizal plants (FmSe, FmMs; Fig. 2; Supplemental Tables S6, S7 and S8). These results point to a more complex regulation in mycorrhizal plants of hormone pathways in response to herbivory.

**Figure 2.**
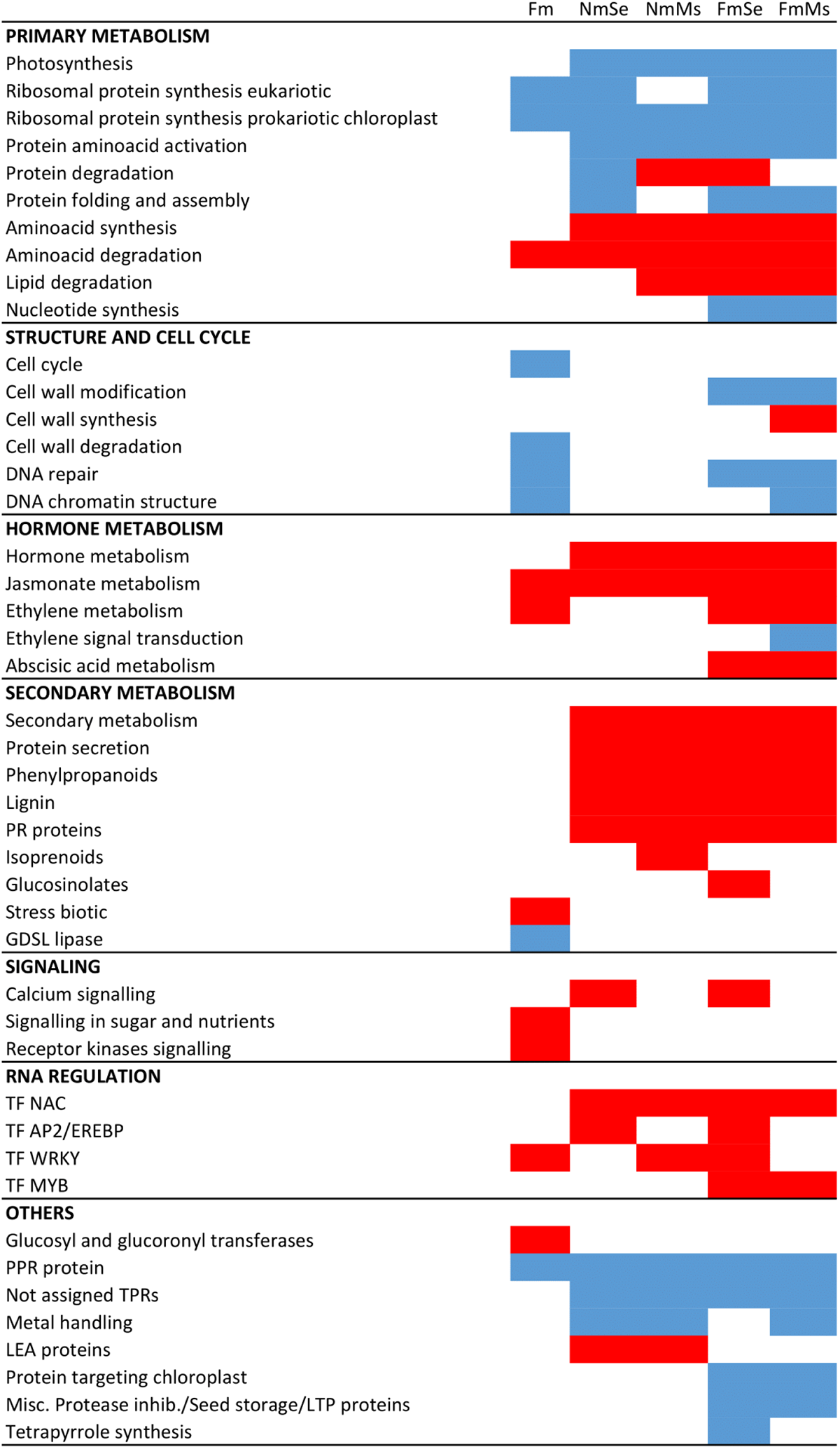
Mycorrhizal colonization impacts plant transcriptional regulation in non-challenged- and herbivory-challenged leaves. Tomato leaves from uninfested non-mycorrhizal (Nm) and *F. mosseae* mycorrhizal (Fm) plants or infested for 24 h with *S. exigua* (NmSe, FmSe) or to *M. sexta* (NmMs, FmMs). Heatmap of enriched gene sets in the different treatments as compared with the non-challenged, non-mycorrhizal control (Nm) according to GSEA (FDR<0.05). Blue and red cells indicate repression and induction of the gene set, respectively.

### 3.3. Mycorrhization primes JA-regulated defense responses upon herbivory

The transcriptomic analysis confirmed the strong induction of the JA signaling pathway upon herbivory (Supplemental Tables S6 and S9). For a more precise analysis of the transcription levels on the JA regulated antiherbivore defenses, we performed targeted analysis of well-characterized JA-regulated anti-herbivory genes: *leucine aminopeptidase A* (*LapA*, Solyc12g010020), *polyphenol oxidase F* (*PPOF*, Solyc08g074620), *threonine deaminase* (*TD*, Solyc09g008670) and *multicystatin* (*MC*, Solyc00g071180). These genes were induced by herbivory in all plants, but the induction was higher in mycorrhizal plants (Fig. 3A), confirming that mycorrhiza leads to primed JA-regulated defenses upon herbivory. We then explored if such changes related to primed JA biosynthesis by analyzing the expression of JA biosynthetic genes *lipoxygenase D* (*LOXD*, Solyc03g122340), *allene oxide synthase 1* (*AOS1*, Solyc04g079730), *allene oxide cyclase* (*AOC*, Solyc02g085730) and *12-oxophytodienoate reductase 3* (*OPR3*, Solyc07g007870). Remarkably, a small, yet significant induction of the JA biosynthesis genes *LOXD* and *AOC* was detected in unchallenged mycorrhizal plants (Fig. 3B). Herbivory induced the expression of JA biosynthesis related genes in both non-mycorrhizal and mycorrhizal plants, but again the induction was overall stronger in mycorrhizal plants, although the primed induction was only significant in the case of *M. sexta* infestation (Fig. 3B). The results confirmed boosted induction upon herbivory of JA synthesis and JA regulated defenses in mycorrhizal plants.

**Figure 3.**
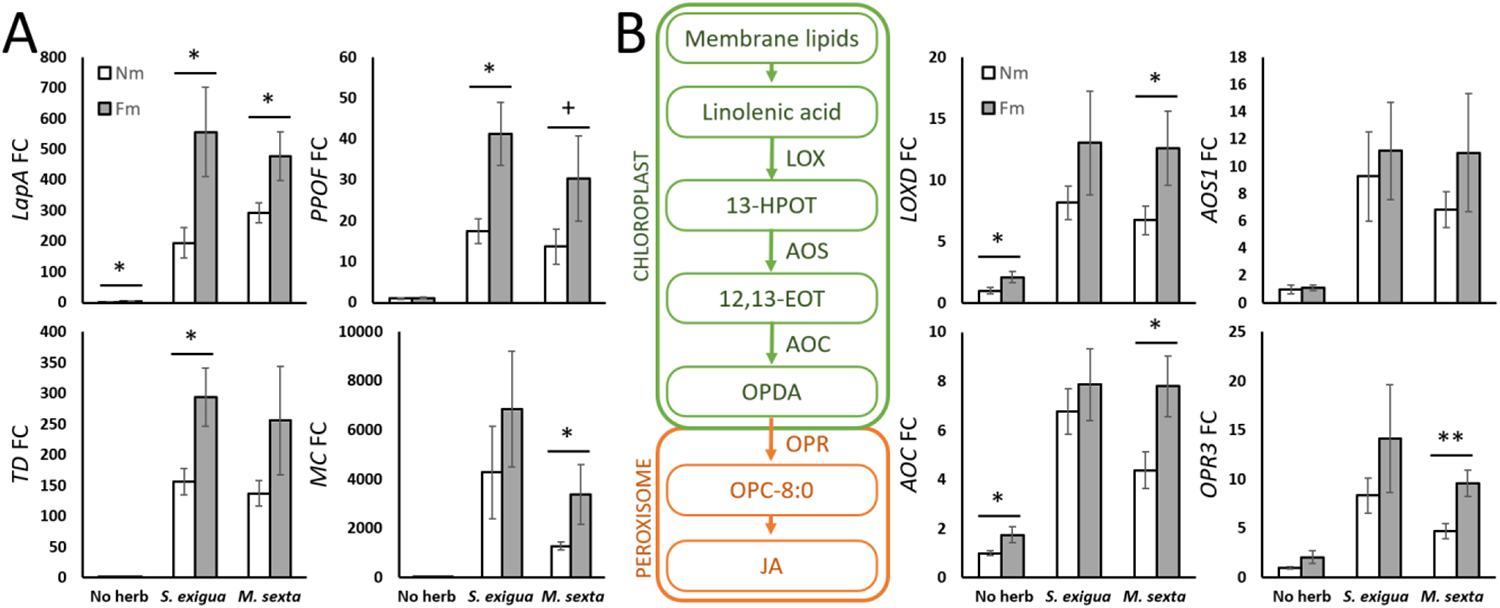
JA-dependent antiherbivory responses and JA biosynthesis upon herbivory are primed in mycorrhizal plants. Tomato leaves from non-mycorrhizal (Nm) and *F. mosseae* mycorrhizal (Fm) plants, uninfested (No herb) or infested for 24 h by *S. exigua* or *M. sexta* (*S. exigua*, *M. sexta*). (A) Relative expression of JA-dependent defense related marker genes: *leucine aminopeptidase A* (*LapA*, Solyc12g010020), *polyphenol oxidase F* (*PPOF*, Solyc08g074620), *threonine deaminase* (*TD*, Solyc09g008670) and *multicystatin* (*MC*, Solyc00g071180); and (B) Relative expression of JA biosynthetic pathway genes: *lypoxigenase D* (*LOXD*, Solyc03g122340), *allene oxide synthase 1* (*AOS1*, Solyc04g079730), *allene oxide cyclase* (*AOC*, Solyc02g085730) and *12-oxophytodienoate reductase 3* (*OPR3*, Solyc07g007870). Expression values were normalized using the reference gene *SlEF*. Data shown are mean ± SEM of 6 biological replicates. Statistical analysis was performed with unpaired t-test analysis between each herbivory treatment. + p<0.1, * p<0.05, ** p<0.01.

### 3.4. Mycorrhization enhances ET metabolism and primes ET biosynthesis and signaling upon herbivory

The RNA-Seq and GSEA revealed differential regulation of ET-related genes in mycorrhizal plants (Supplemental Tables S5, S7 and S10). To more precisely quantify the transcriptional differences related to ET signaling, we performed a targeted analysis of transcriptional regulation of ET biosynthesis and signaling marker genes (Fig. 4A). Higher basal levels in mycorrhizal plants were confirmed for the ET biosynthetic genes *ACC synthase* (*ACS6*; Solyc08g008100) and *ACC oxidases* (*ACO1* and *ACOlike4*; Solyc07g049530 and Solyc04g007980), and for an ethylene responsive factor (ERF; Solyc02g070040). These genes were also upregulated in response to both herbivores. Remarkably, as in the case of JA-genes genes (Fig. 3) the induction by herbivory of the ET-related genes was generally higher in mycorrhizal plants (Fig. 4A). This primed response was stronger in mycorrhizal plants challenged with *M. sexta* (Fig. 4A), where more JA-regulated mycorrhiza-related changes were also seen (Fig. 3B). We then quantified ET emission (in a new set of plants) to evaluate the relevance of the observed transcriptional upregulation of ET biosynthesis in mycorrhizal plants. Leaves of *F. mosseae* plants emit significantly more ET than those from non-mycorrhizal plants (Fig. 4B). Herbivory treatments increased ET production in both mycorrhizal and non-mycorrhizal plants, but mycorrhizal plants had slightly higher levels in *M. sexta* infested leaves (p<0.07). The results confirm a differential regulation of ET biosynthesis in mycorrhizal plants.

**Figure 4.**
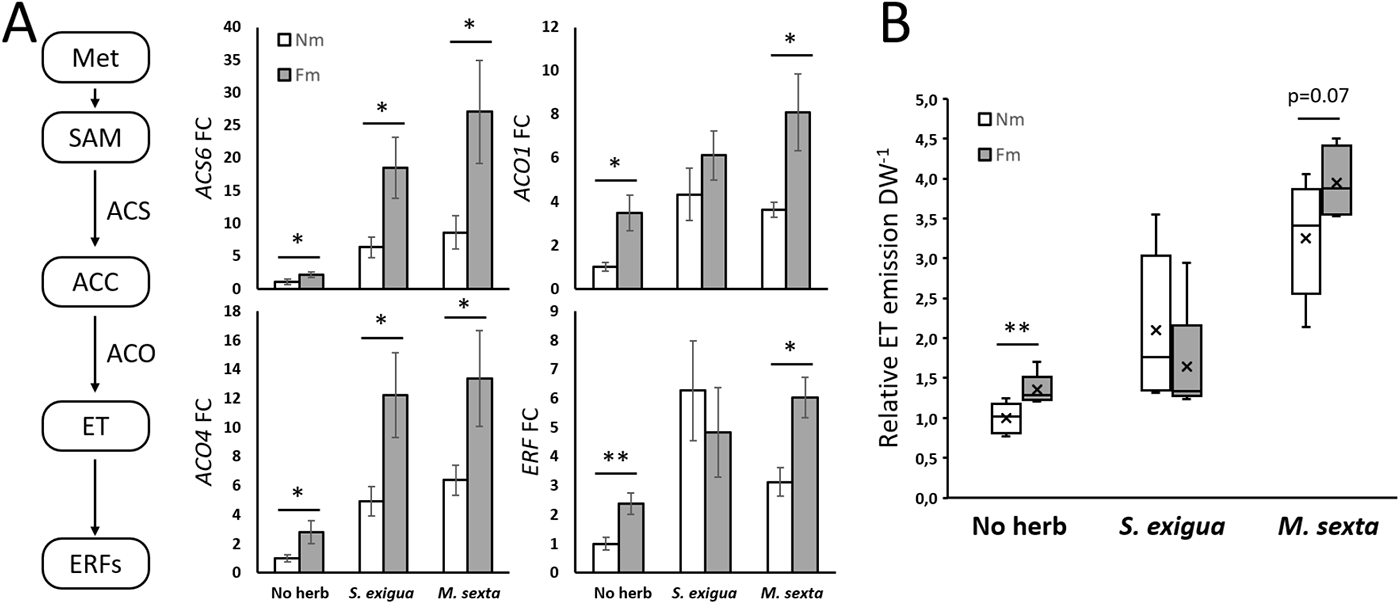
Mycorrhizal symbiosis primes ethylene biosynthesis and signaling. Tomato leaves from non-mycorrhizal (Nm) and *F. mosseae* mycorrhizal (Fm) plants, uninfested (No herb) or infested for 24 h by *S. exigua* or *M. sexta* (*S. exigua*, *M. sexta*). (A) Relative expression of ET biosynthesis genes *1-aminocyclopropane-1-carboxylic acid (ACC) synthase 6* (*ACS6*, Solyc08g008100), *ACC oxidase 1* (*ACO1*, Solyc07g049530) and *ACC oxidase 4-like* (*ACO4-like*, Solyc04g007980) and an *ET responsive factor* (*ERF*, Solyc02g070040). (B) Boxplots show relative ET emission normalized to leaflet dry weight (DW). Single tomato leaflets of non-mycorrhizal plants (Nm) and mycorrhizal plants with *F. mosseae* (Fm) were challenged with *S. exigua* or *M. sexta* for 18 h inside 20 mL glass vials. 1 mL of every sample was withdrawn from the vial and the area of the ethylene peak was analyzed by gas chromatography. (A) Expression values were normalized using the reference gene *SlEF*. Data shown are mean ± SEM of 6 (A) or 5 (B) biological replicates. Statistical analysis was performed with unpaired t-test analysis between each herbivory treatment. + p<0.1, * p<0.05, ** p<0.01.

### 3.5. Physical interaction networks supports a connection of ET signaling with JA biosynthesis

To explore the potential interaction between the ET and JA signaling pathways in the differential response of mycorrhizal tomato plants to *S. exigua* and *M. sexta*, the gene expression data were plotted into a physical interaction network, constructed by merging dispersed resources on metabolic pathways, protein-protein interactions, protein-DNA interactions and small RNA-transcripts interactions (Ramšak *et al.,* 2018). We next extracted a subnetwork of genes differentially expressed when comparing leaves of mycorrhizal and non-mycorrhizal plants and their direct interactors. This network was further explored by extraction of the shortest paths between the nodes with main function in ET signaling (EIN3 and EIN3-like nodes) (left side of the network) and the ones participating in JA biosynthesis (right side of the network) (Fig. 5). The results show a broad activation of the JA and ET pathway genes in mycorrhizal plants in the absence of herbivory (Fig. 5A). In plants subjected to herbivory (Fig. 5 B and C), the symbiosis also had an impact on the two pathways, pointing to an interconnected regulation between both hormones that may lead to the differential regulation of the plant responses to herbivory in mycorrhizal plants. Particularly, the expression of *WRKY40* was induced in all three comparisons, with higher levels in mycorrhizal plants regardless the herbivory status. In addition, ET responsive factors *ERF16* (ortholog of the Arabidopsis *ORA47*), *CBF1* and *CBF2,* proposed to act on the JA biosynthetic genes, were differentially regulated in mycorrhizal plants in all three comparisons (Fig. 5). The network results, based on previous experimental data on connections between the different elements, illustrate the interconnected differential regulation of ET and JA signaling elements in mycorrhizal plants. They also pinpoint key regulatory elements potentially mediating the regulatory role of ET on the JA both pathway leading to MIR.

**Figure 5.**
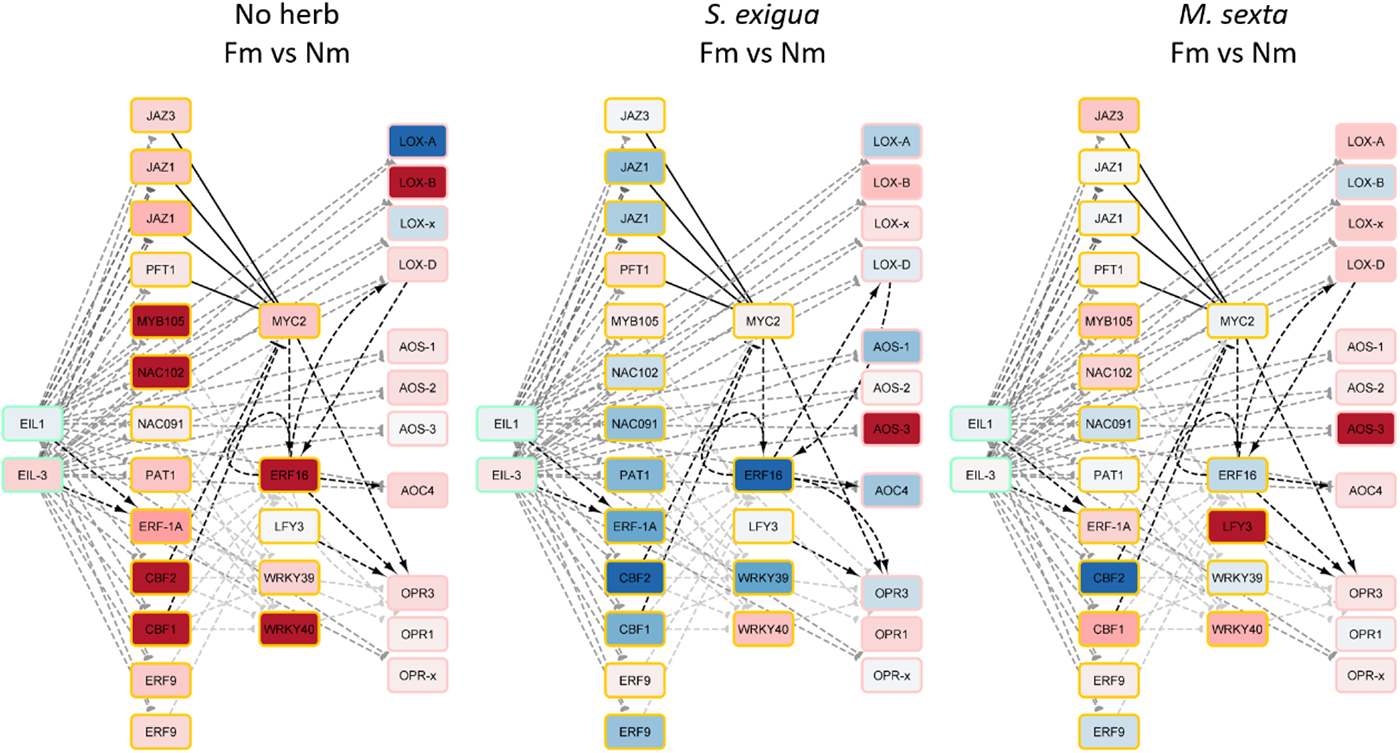
Network analysis visualization for mycorrhizal effects on the JA and ET pathways in response to herbivory. Tomato leaves from non-mycorrhizal (Nm) and *F. mosseae* mycorrhizal (Fm) plants, uninfested (No herb) or infested for 24 h by *S. exigua* or *M. sexta* (*S. exigua*, *M. sexta*). Insight into the network of physical interactions between the signaling network components. Nodes represent tomato protein-coding genes and the node color shows regulation after the applied treatment (red = up-regulation, blue = down-regulation). Connection type is shown as different line types (dashed = transcriptional regulation; solid = binding or synthesis); line arrows show the mode of action (arrow = activation; T = inhibition; half-circle = unknown).

### 3.6. Ethylene signaling is essential for the display of MIR against herbivory

As the network analysis identified potential direct connections of ET signaling with JA biosynthesis and signaling, we hypothesized that ET signaling regulates the boosted JA-related antiherbivore responses triggered by the AM symbiosis. To test this, we performed an experiment using tomato lines impaired in ET synthesis (ACD, expressing the *Pseudomonas* ACC deaminase that cleaves the ET precursor ACC; Klee *et al.,* 1991) or ET perception (Nr: Never ripe, mutant in the ET receptor ETR3, Wilkinson *et al.,* 1995), both in a UC82B background. We first confirmed that mycorrhizal colonization was well established in all lines, with no significant differences between the genotypes (Supplemental Fig. S3A). There was no evident effect of the symbiosis on plant biomass in any of the plant genotypes (Supplemental Fig. S3B). We then analyzed ET emission in the different lines. ET was induced upon herbivory in the wild-type, up to 2.5-fold after 3h of feeding by *M. sexta* larvae (Supplemental Fig. S4). Herbivore-induced ET accumulation was significantly higher in mycorrhizal plants, confirming the primed accumulation of ET in response to herbivory. ET production was almost abolished in the ACD mutant regardless of the treatment, and while ET production still increased in response to herbivory in the ET insensitive mutant Nr, the primed response by mycorrhiza was lost (Supplemental Fig. S4).

We next explored the role of ET in MIR by evaluating MIR in the ET-deficient lines. There was a slight, non-significant, reduction in larval survival (Fig. 6A), and a reduction in the total number of pupae in the ET deficient lines as compared to the wild-type (Fig. S5), which aligns with the traditionally described negative crosstalk between ET and JA-mediated herbivory defenses (Verhage *et al*., 2011; Kazan & Manners, 2013; Zhang *et al.,* 2018). This is in agreement with a negative role of ET on basal resistance to herbivory. However, when analyzing the effect of mycorrhizal colonization on plant resistance, mycorrhizal colonization led to a significant increase in mortality of *S. exigua* larvae compared to non-mycorrhizal plants in wild-type (UC82B) plants, but this increased mortality and impaired pupation was abolished in both ET deficient lines, ACD and Nr (Fig. 6A, Fig. S5). Thus, ET plays a positive role in the induced resistance.

Regarding the performance of the specialist *M. sexta*, larval mortality was almost null in all the treatments (Supplemental Fig. S6), this low mortality probably associated to its high adaptation for Solanaceae hosts. Still, larval biomass determination revealed a significant negative effect of the mycorrhizal symbiosis on *M. sexta* performance in the wild-type, as the larvae reared on Fm plants had lower biomass compared to those reared on Nm plants (Fig. 6B). This reduction was also lost in the ET-impaired lines (Fig. 6B). Thus, the bioassays with both herbivores demonstrate that ET signaling is required for MIR.

**Figure 6.**
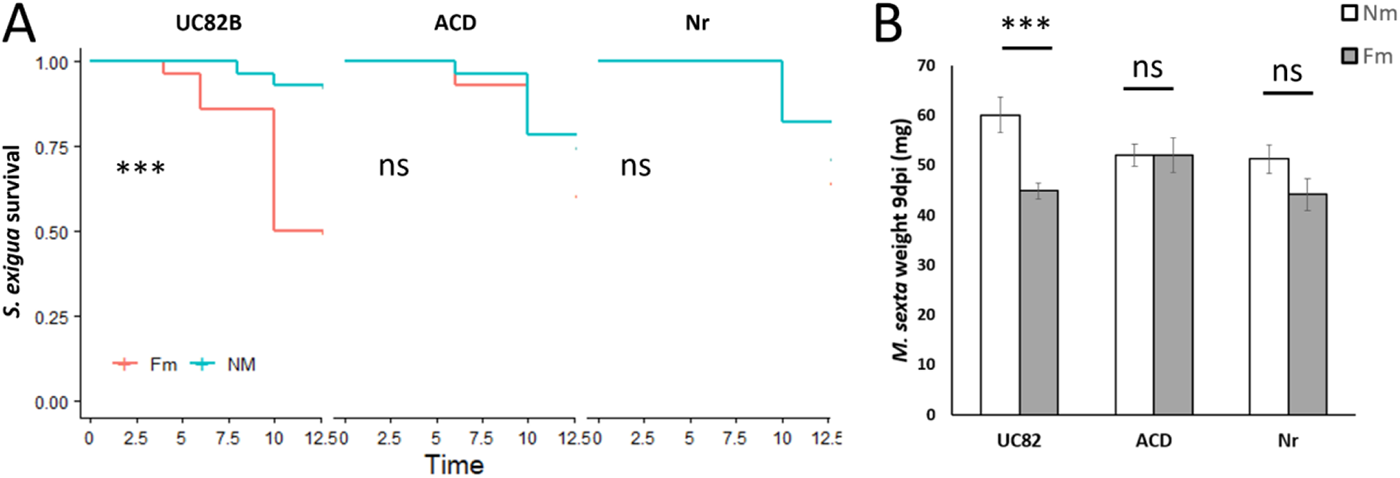
ET deficient lines do not display MIR. (A) Larval survival was monitored at every 2-3 days for *S. exigua* larvae, and (B) *M. sexta* larval biomass determined 9 dpi after feeding on tomato plants of non-mycorrhizal (Nm) and *F. mosseae* mycorrhizal (Fm) plants in the wild-type (UC82B) or ET deficient lines (ACD, Nr). We placed (A) 4 3^rd^ *S. exigua* larvae or (B) 3 neonate *M. sexta* larvae on the plant’s first true leaf and let them feed inside an entomological bag of (A) 7 plants (n=28 larvae) and (B) 10 plants (n=30 larvae) per treatment. Before they had consumed the whole leaf, we moved them to the next consecutive leaf. Statistical analysis was performed with (A) differences between curves estimated with a logrank (Mantel-Cox) test and (B) unpaired t-test analysis between each genotype. *** p<0.001.

### 3.7. Ethylene as a positive regulator of MIR-related priming of JA biosynthesis

Finally, to explore the mechanism of MIR regulation by ET, and according to the direct connection between ET and JA signaling pointed by the network analysis (Fig. 5), we evaluated JA-regulated antiherbivory defenses in the ET-deficient lines ACD and Nr. We focused on the interaction with *M. sexta* because it overall showed the greatest transcriptional changes in mycorrhizal plants as compared with non-mycorrhizal ones (Figs. 2, 3 and 4). We first addressed if the primed JA regulated defenses to herbivory in mycorrhizal plants require functional ET signaling. In wild-type herbivory-challenged plants, the expression of *LapA*, and the corresponding enzymatic activity, were boosted in mycorrhizal plants compared to non-mycorrhizal ones. This primed response in mycorrhizal plants was completely abolished in both ET deficient lines (Fig. 7A), supporting that ET is required for the priming of JA-regulated defenses associated to MIR.

We next aimed to further disentangle how ET regulates JA-dependent responses during MIR. With this aim, we explored the role of ET signaling in the mycorrhiza-mediated priming of JA biosynthesis. We first analyzed the expression of genes pointed by the network analysis (Fig. 5) as candidates to mediate ET regulation of JA biosynthesis upon herbivory. We evaluated the expression levels of the transcription factors *WRKY39* and *WRKY40* (Solyc03g116890 and Solyc06g068460), *LFY3* (Solyc03g118160) *MYC2* (Solyc08g076930), and the APETALA2/Ethylene-response factors (AP2/ERF) *ERF16* and *ERF15* (Solyc12g009240 and Solyc06g054630; *ORA47* orthologs), and *CBF1* and *CBF2* (Solyc03g026280 and Solyc03g124110) in the wild-type and the ET-deficient lines under herbivory. Differential expression in wild-type mycorrhizal plants was found for *WRKY39*, *WRKY40*, *CBF1*, *CBF2,* and *MYC2* (Fig. 7B; Supplemental Fig. S7). Again, these mycorrhiza-associated changes were lost in the ET-deficient lines (Fig. 7B; Supplemental Fig. S7). Last, we analyzed the expression of the main JA biosynthetic genes (Fig. 7C). In wild-type challenged plants, mycorrhization led to boosted expression of *LOXD* and *OPR3,* and a similar - although not significant-trend in *AOS1* (Fig. 7C), whereas *AOC* expression was not altered (Supplemental Fig. S7). Finally, we quantified by UPLC-MS the metabolites OPDA, JA, and JA-Ile, and in agreement with the increased expression of the biosynthesis genes, we found a higher accumulation of JA and JA-Ile in wild-type challenged mycorrhizal plants, as compared to non-mycorrhizal plants (Fig. 7D). This boosted expression of JA biosynthetic genes and in jasmonates accumulation triggered by mycorrhiza was completely lost in the ET impaired lines (Fig. 7, C and D). Our results confirm that ET is required for the mycorrhizal priming of JA biosynthesis in herbivory challenged plants leading to MIR.

**Figure 7.**
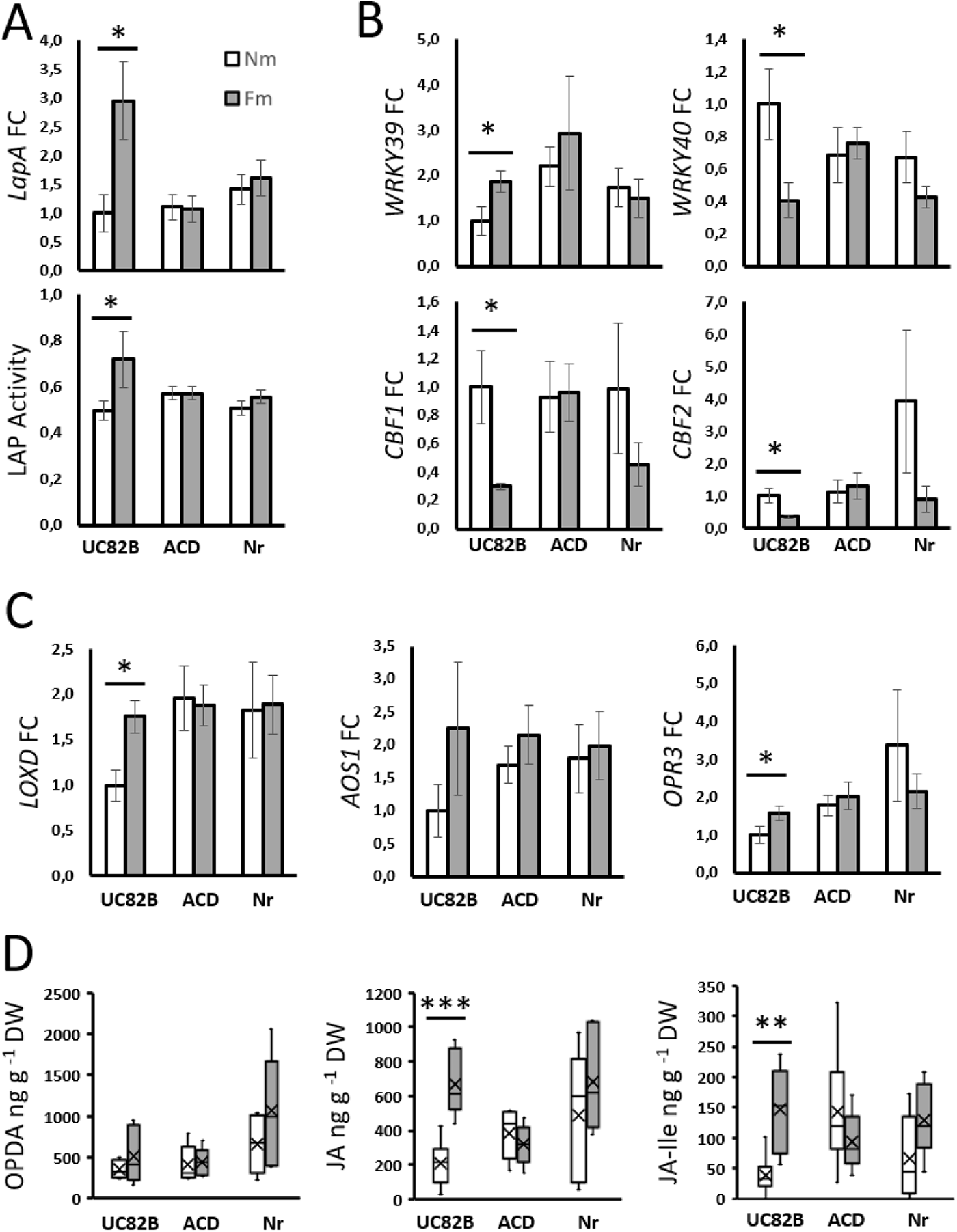
ET is required for the primed JA burst in herbivory-challenged mycorrhizal tomato plants. Tomato plants of non-mycorrhizal (Nm) and *F. mosseae* mycorrhizal (Fm) plants in the wild-type genotype (UC82B) or ET deficient lines (ACD, Nr) subjected to *M. sexta* herbivory. 3 larvae were added per plant, and newly infested leaves were harvested 24 h after infestation. (A) Relative expression of the JA-dependent defense related marker gene *leucine aminopeptidase A* (*LapA*, Solyc12g010020) and the corresponding enzymatic activity (LAP). (B) Relative expression of JA-ET related transcription factor genes: *WRKY39* (Solyc03g116890), *WRKY40* (Solyc06g068460), *CBF1* (Solyc03g026280) and *CBF2* (Solyc03g124110). (C) Relative expression of JA biosynthetic pathway genes: *lipoxygenase D* (*LOXD*, Solyc03g122340), *allene oxide synthase 1* (*AOS1*, Solyc04g079730) and *12-oxophytodienoate reductase 3* (*OPR3*, Solyc07g007870). (D) Levels of different oxylipins/JA metabolites (OPDA, JA, and JA-Ile) in the challenged leaves determined by UPLC-MS (A, B, C). Data represent mean ± SEM of 6 biological replicates. Expression values were normalized using the reference gene *SlEF*. (D) Boxplots of 6 biological replicates normalized to plant dry weight. Statistical analysis was performed with unpaired t-test analysis between each herbivory treatment. * p<0.05, ** p<0.01, *** p<0.001.

## 4. Discussion

Mycorrhiza-induced resistance (MIR) has been described in different plant species, mostly against necrotrophic pathogens and leaf-chewing insects (Hartley & Gange, 2008; Song *et al.,* 2013; Roger *et al.,* 2013; Nair *et al.,* 2015; Sanchez-Bel *et al.,* 2016; He *et al.,* 2017; Sanmartín *et al.,* 2020b,a; Rivero *et al.,* 2021). The susceptibility of these aggressors to JA-regulated plant defense responses suggested the involvement of JA signaling in MIR (Pozo & Azcón-Aguilar, 2007). This hypothesis was supported by results showing transcriptional regulation of JA dependent defenses associated to MIR, and the use of JA-deficient lines confirmed the central role of JA as a main regulator of MIR (Song *et al*., 2013; Mora-Romero *et al*., 2014). Further studies including omics approaches illustrated the complexity of MIR associated responses, suggesting the existence of additional regulatory elements in the process (Sanmartín *et al*., 2020b; Rivero *et al*., 2021), but the molecular regulation of defense priming in MIR remains uncovered. Here we followed an untargeted approach to identify novel elements in the regulation of MIR against chewing herbivores in tomato. Our analysis revealed primed JA-regulated antiherbivore responses in mycorrhizal plants, and pinpointed ET as a potential key regulatory element of the primed response. By combining transcriptomic, enzymatic and metabolomic analyses, herbivore bioassays and a genetic approach using ET deficient lines we were able to demonstrate that ET signaling plays a central role in MIR.

Mycorrhizal colonization has a strong impact on root transcriptome and metabolome profiles, as shown in different plants including tomato (López-Ráez *et al*., 2010; Rivero *et al*., 2015, 2018; Vangelisti *et al*., 2018; Hsieh *et al*., 2022; Jing *et al*., 2022), but usually generate only minor changes in the leaves in the absence of challenge (Song *et al.,* 2013; Schweiger *et al.,* 2014; Adolfsson *et al*., 2017; Rivero *et al*., 2021). In our full transcriptome analysis, we also found a low number of differentially expressed genes in mycorrhizal plants in the absence of challenge, but more differences were observed upon herbivory, fitting with a primed plant response to the aggressor. Nevertheless, while few significant DEGs were identified in mycorrhizal plants in the absence of challenge, gene set enrichment analyses revealed that the symbiosis impacts diverse functional categories that could contribute to their primed state. For example, mycorrhizal symbiosis triggered changes related to transcriptional regulation, in histone and chromatin compaction as well as in the abundance of transcription factors and receptor kinases. These elements have been proposed to underlie the primed state in preconditioned plants (Conrath *et al*., 2015). In addition, the categories corresponding to JA and ET metabolism were also upregulated by mycorrhiza at the basal level, pointing to differential hormone homeostasis in these plants. All these changes are consistent with the premise that defensive priming generates few basal changes in the organism, but in key regulatory aspects, that may allow a boosted response upon challenge (Martinez-Medina *et al*., 2016; Mauch-Mani *et al*., 2017). Indeed, upon herbivory, differences between mycorrhizal and non-mycorrhizal plants were more pronounced, supporting a primed response to the attack.

Herbivory led to a strong transcriptional reprogramming in the plant, regardless of its mycorrhizal status. The transcriptomic analysis evidenced a core of common plant responses to both herbivores, including the upregulation of plant secondary metabolism for defense, and the reorganization of primary metabolism. Nevertheless, we observed some variations in the response to the herbivores, as some categories were altered in the interaction with the generalist *S. exigua* but not with the specialist *M. sexta*, which could be related to their degree of specialization (Ali & Agrawal, 2012), as specialists try to circumvent plant defense activation. Certainly, both herbivores activated the JA signaling pathway, as it is the central pathway of resistance against chewing herbivores (Wasternack & Hause, 2013; Erb & Reymond, 2019).

While these responses to herbivory were overall common in non-mycorrhizal and mycorrhizal plants, more changes were detected in mycorrhizas, with some exclusive changes in categories related to hormone metabolism, cell wall modifications, transcription factors, and protease inhibitors. Particularly, the induction of JA regulated responses was boosted in mycorrhizal plants, as they displayed a stronger transcriptional activation of antiherbivory defenses, for example of genes coding for the well characterized defensive Leucyl aminopeptidase A (LAPA), multicystatin (MC), threonine deaminase (TD) and polyphenoloxidases (PPOs) (Green & Ryan, 1972; Felton *et al.,* 1989; Chen *et al.,* 2005; Erb & Reymond, 2019).

Remarkably, besides the activation of JA responses, ABA and ET pathways were also differentially regulated in mycorrhizal plants. Such changes in additional signaling pathways could contribute to a more complex regulation of antiherbivore responses. Indeed, hormone crosstalk is key in shaping final defense responses in plants (Erb *et al*., 2012; Berens *et al*., 2017; Aerts *et al*., 2021). ET is an essential regulator of plant defenses, playing a complex modulatory role in the overall hormonal crosstalk shaping plant defense responses to specific challenges (Broekgaarden *et al*., 2015). While it is well documented that herbivory damage induces ET emission in plants (Winz & Baldwin, 2001; de Vos *et al*., 2005), ET signaling is believed to play a negative role in resistance to herbivores. For example, in Arabidopsis, the ABA/JA synergy orchestrates an effective defense response to chewing insects, that is finetuned through antagonism with the ET-regulated pathway (Bodenhausen & Reymond, 2007; Verhage *et al*., 2011). Antagonism between ET and ABA/JA pathways also impairs nicotine synthesis against *M. sexta* larvae in *Nicotiana attenuata,* resulting in diversified defensive responses (von Dahl & Baldwin, 2007; Onkokesung *et al*., 2010). Not surprisingly, attackers can modulate ET signaling to alter hormone crosstalk for their benefit (Vogel *et al*., 2007), and for example, elicitors in *S. exigua* saliva manipulate ET signaling to suppress JA responses in *Medicago* (Paudel & Bede, 2015). On the other hand, ET/JA signaling are required for phenolamide accumulation in response to herbivory (Figon *et al.,* 2021; Li *et al.,* 2020; Li *et al.,* 2018; Wang *et al.,* 2020), and recent studies points to potential synergy of JA and Et also in resistance to herbivory (Hu *et al.,* 2021). Thus, experimental evidence illustrate the complex regulatory role of ET in shaping defensive responses, probably finetuning defenses through positive and negative interactions with other pathways depending on timing and hormone doses.

Our analyses revealed that mycorrhizal symbiosis enhanced ET synthesis in leaves under basal conditions, and these increased ET levels are further amplified upon herbivory. We hypothesized that this differential regulation of the ET pathway in mycorrhizal plants is essential for MIR against herbivory. The relevance of JA-ET interplay in microbe-induced plant resistance has been reported against necrotrophic pathogens, mostly in the Arabidopsis-*Pseudomonas* WCS417r model system (van Loon *et al*., 2006; Pieterse *et al*., 2014). However, the involvement of ET in induced resistance against herbivores has been only proposed in the Arabidopsis-Pseudomonas WCS417r induced resistance against *Mamestra brassicae* (Pangesti et al. 2016). In our study, we show that the tomato ET deficient lines ACD (deficient in ET production) and Never ripe (deficient in ET signaling) were unable to display MIR against the two herbivores tested, despite similar mycorrhizal colonization levels. Regarding plant defense responses, wild-type mycorrhizal plants showed primed expression and activity levels of LapA upon herbivory. LapA is considered a marker of JA regulated antiherbivore defenses, but in addition, it acts as a chaperone regulating late JA and wound response (Fowler *et al*., 2009). Accordingly, the changes in LapA levels may have further consequences in the plant defensive response through differential post-transcriptional modifications. Remarkably, the primed LapA activity in mycorrhizal plants, and the enhanced resistance to the herbivores, was lost in the ET-deficient mutants, confirming that ET is required for mycorrhizal priming of JA regulated antiherbivore defenses and MIR.

Finally, the transcriptome and network analyses results suggested that the ET effect may act upstream of JA signaling, likely targeting JA biosynthesis. The effect of ET on JA biosynthesis in response to wounding was only modestly studied earlier. Previous studies demonstrated that inhibition of ET signaling partially inhibited JA synthesis (O’Donnell et al. 1996), and ORA47, an APETALA2/Ethylene-response factor (AP2/ERF) were shown to positively regulate OPDA biosynthesis by binding to the promoters of most JA biosynthetic genes (Pauwels *et al*., 2008; Chen *et al*., 2016; Hickman *et al*., 2017). Thus, considering the described antagonism and synergism between ET and JA, evidences suggest a modulatory role of ET that combines both positive and negative effects on the JA pathway. Timing in the sequential induction of phytohormones seems to be essential for the crosstalk outcome: In late stages of herbivory attack, ET may be antagonistic to JA-defenses, but ET seems to be required for the JA-burst upon challenge. Our network analysis pointed to some regulatory elements as candidates to mediate the positive impact of ET signaling on JA biosynthesis, and differential regulation in mycorrhizal plants was confirmed for the ET responsive factors *WRKY39*, *WRKY40, CBF1 and CBF2*. The follow-up analyses on the ET deficient lines revealed that their differential regulation, the boosted expression of the JA-biosynthesis genes *LOXD* and *OPR3*, as well as the enhanced accumulation of JA and its bioactive conjugate JA-Ile observed in mycorrhizal wild-type plants was lost in the ET-deficient lines. Thus, our study demonstrates that the regulation of ET signaling in mycorrhizal plants is required for the primed accumulation of JA upon herbivory in mycorrhizal plants. The results agree with recent studies showing that in tomato, *ORA47* orthologs *ERF15* and *ERF16* are responsible for the quick transcriptional activation of JA synthesis upon herbivory by inducing *LOXD*, *AOC* and *OPR3* transcription (Hu *et al.,* 2021).

In conclusion, our study provides comprehensive evidence that ET signaling plays a critical role in the primed defense responses that lead to enhanced resistance in mycorrhizal plants. The differential regulation of ET biosynthesis in mycorrhizal plants leads to a boosted JA burst upon herbivore attack, and the primed activation of downstream JA-dependent responses. Our results contribute to elucidating the complex hormone crosstalk in shaping plant defense priming in mycorrhizal plants, with ET acting as a pivotal regulator of JA defenses upon herbivory. Understanding the regulation of plant defense priming by beneficial microbes paves the way for improving biotechnological applications of microbial inoculants in sustainable crop protection.

## Acknowledgements

We acknowledge Javier Palenzuela (EEZ-CSIC, Spain) for kindly providing the *F. mosseae* inocula. Serveis Centrals d’Instrumentació Científica (SCIC) at the Universitat Jaume I for the hormone analysis. Iván Fernandez, Juan García, Estefanía Berrio, Andrea Ramos and Luis España for their valuable technical support. Nicole van Dam for support and helpful discussions. Salvador Herrero (Universitat de València, Spain) and Felix Tiburcy and Harald Mikoleit (Universität Osnabrueck, Germany) for kindly providing us *S. exigua* and *M. sexta* eggs. Harry Klee (Florida, USA) for kindly providing us UC82B, ACD and Nr seeds.

## Competing interests

The authors declare that the research was conducted in the absence of any commercial or financial relationships that could be construed as a potential conflict of interest.

## Author contributions

JL, JR, AMM and MJP contributed to the conception and design of the study. JR and AMM performed the transcriptional bioassay. JL, ŽR, MP and KG performed the transcriptional analysis. MK constructed the tomato-specific miRNA-transcript network. JL performed the functional bioassays and the corresponding molecular analysis. VF performed metabolomics analyses. JL performed statistical analyses. JL and MJP wrote the manuscript and AMM, KG, MP, ŽR, MK, JLR and VF revised it.

## Data Availability

RNA-Seq data is openly available at National Center for Biotechnology Information (NCBI) Sequence Read Archive (SRA) (https: //www.ncbi.nlm.nih.gov/sra) under the SRA accession SRP458983. The raw data that support the findings of this study are available from the corresponding author upon request.

## Funding

This research has been funded by projects RTI2018-094350-B-C31 and PID2021-124813OB-C31 from the Spanish National R&D Plan of the Ministry of Science, Innovation and Universities (MICIU), and the European Regional Development Fund (ERDF) ‘a way a making Europe’. We acknowledge ideas exchange under COST Action FA1405.

## Supporting information

**Supplemental Fig. S1.** Heatmap of S. exigua treatment (NmSe) changes on enriched gene sets compared with M. sexta treatment (NmMs)

**Supplemental Fig. S2**. Heatmap of mycorrhizal herbivory treatments changes on enriched gene sets compared with their non-mycorrhizal herbivory controls

**Supplemental Fig. S3**. Mycorrhizal root colonization and shoot biomass of wt and ET deficient lines

**Supplemental Fig. S4**. Relative ET emission of non-mycorrhizal and mycorrhizal wt and ET deficient lines

**Supplemental Fig. S5**. S. exigua pupation in non-mycorrhizal and mycorrhizal wt and ET deficient lines

**Supplemental Fig. S6**. M. sexta mortality in non-mycorrhizal and mycorrhizal wt and ET deficient lines

**Supplemental Fig. S7**. Relative expression of JA-ET related transcription factor genes in wt and ET deficient lines after 24h of M. sexta herbivory

**Supplemental Table S1**. GSEA Manually organized functional supergroups from enriched gene sets

**Supplemental Table S2**. Primers used for qPCR

**Supplemental Table S3**. RNA-seq DEGs overview

**Supplemental Table S4**. Total processed RNA-Seq table

**Supplemental Table S5**. RNA-seq Fm vs Nm DEGs

**Supplemental Table S6**. GSEA JA

**Supplemental Table S7**. GSEA ET

**Supplemental Table S8**. GSEA ABA

**Supplemental Table S9**. RNA-seq read counts JA

**Supplemental Table S10**. RNA-seq read counts ET

